# Astrocyte senescence in an Alzheimer’s disease mouse model is mediated by TGF-β1 and results in neurotoxicity

**DOI:** 10.1101/700013

**Authors:** Shoshik Amram, Tal Iram, Ekaterina Lazdon, Robert Vassar, Ittai Ben-Porath, Dan Frenkel

## Abstract

Alterations in astrocyte function such as a pro-inflammatory phenotype are associated with Alzheimer’s disease (AD). We had shown impairments in the ability of aged astrocytes isolated from 5xFAD mice to clear and uptake amyloid-β (Aβ) as well as to support neuronal growth. Senescent cells accumulate with age and exhibit a senescence-associated secretory phenotype, which includes secretion of pro-inflammatory cytokines. In this study, we predicted that with age, astrocytes in 5xFAD mice would exhibit a cellular senescence phenotype that could promote neurodegeneration. We found an age-dependent increase in senescent astrocytes adjacent to Aβ plaques in 5xFAD mice. Inhibition of nuclear factor kappa-light-chain-enhancer of activated B cells reduced interelukin-6 secretion by senescent astrocytes and resulted in improved neuronal support. Moreover, senescent astrocytes exhibited an increase in the induction of the TGF-β1-SMAD2/3 pathway, and inhibition of this pathway resulted in a reduction of cellular senescence. We also discovered that soluble Aβ42 induced astrocyte senescence in young naïve mice in a SMAD2/3-dependent manner. Our results suggest an important role of astrocyte senescence in AD and its role in mediating the neurotoxicity properties of astrocytes in AD and related neurodegenerative diseases.

## INTRODUCTION

Cellular senescence (CS) is a process in which cells cease to divide and undergo morphological and functional alterations (1). Cyclin-dependent kinase inhibitors, such as p15^ink4b^, p16^ink4a^, and p21^Cip1^, enforce cell growth arrest by activating the retinoblastoma protein (2), and they serve as the most common markers of CS. Senescent cells are also characterized by a pro-inflammatory phenotype, the senescence-associated secretory phenotype (SASP) (3). The SASP includes a secretion of pro-inflammatory cytokines, such as interleukin (IL)-6, chemokines, such as IL-8, growth factors, such as transforming growth factor β (TGF-β), and proteases (3). With age, there is an increase in cells expressing CS markers in several tissues (4), including the brain (5).

Senescent cells have been shown to disrupt normal tissue organization (6). For example, progeria syndrome is associated with accelerated aging in children (7), and clearance of senescent cells in a progeria syndrome mouse model resulted in a rejuvenation of aged tissues, such as adipose tissue, skeletal muscle, and the eye (7). Furthermore, a clearance of senescent cells in wild-type (WT) mice resulted in an extended life span and mitigated age-related impairments of several tissues (8). Recent studies have shown that clearance of senescent glial cells in a tauopathy mouse model attenuated brain disease pathology (9).

Alzheimer’s disease (AD) is a neurodegenrative disease associated with dementia in the elderly and, although the mechanisms involved are not well understood, the greatest risk factor for AD is aging. Astrocyte activation occurs in early stages of disease pathogenesis, and it was detected even before amyloid beta (Aβ) plaque deposition (10). We recently reported that astrocytes from an aged AD mouse model failed to support neurons and to mediate clearance of Aβ (11). Others suggested that astrocytes in the aging brain express characteristics of CS, which includes a pro-inflammatory profile (12). Moreover, several studies have shown that senescent astrocytes accumulate in greater amounts in the brain with aging (13, 14) and, specifically, in patients with AD compared to age-matched controls (14). The mechanisms underlying astrocyte senescence (AS) and the effects of AS on disease course, however, remain unclear.

In this study, we aimed to identify pathways that are affiliated with AS during the progression of amyloid pathology in 5xFAD mice and to assess potential therapeutic interventions to reduce astrocyte neurotoxicity. Our results revealed that astrocytes adjacent to plaques expressed markers of CS in an age-dependent manner, and that senescent astrocytes showed impairment in supporting neurons, which was reversible by targeting specific pathway.

## MATERIALS AND METHODS

### Experimental Procedures

#### Mice

Heterozygous TGF-β1 mice were acquired from Tony Wyss–Coray’s laboratory (15) and maintained on an inbred C57BL/6 mouse genetic background (Jackson Laboratory). 5xFAD mice that express 5 familial AD (FAD) mutations in amyloid precursor protein and presenilin 1 were acquired from Robert Vassar’s laboratory (16) and maintained on an inbred C57BL/6 mouse genetic background (Jackson Laboratory). All experiments were carried out on male mice. The care and experimental use of all animals were performed in accordance with the Tel Aviv University guidelines and were approved by the University’s animal care committee.

#### Immunohistochemistry

Both the frozen brain slices and the coverslips were fixed using 4% paraformaldehyde diluted in phosphate buffered saline (PBS) for 10 minutes, followed by 5-minute washes with PBS that were repeated 3 times. To decrease background staining, we blocked the slices with blocking medium (8% horse serum, 0.3% Triton, 1 g/100 mL bovine serum albumin, 88.7% PBS, 0.02% sodium azide) for 1 hour. Samples were stained with the following antibodies: mouse monoclonal 6E10 (1:500, Biolegend, #sig39320), rat anti-GFAP (1:500, Invitrogen, #13-0300), rabbit anti-p15 (1:50, Assay Biotechnology, # C0287), rabbit anti-laminb1 (1:200, abcam, # 16048), or mouse anti-β-tubulin antibody (1:1000, Cell Signaling, #2146). Next, the samples were washed 3 times with PBS and stained with secondary antibodies (Alexa Fluor,1:500; 594 goat anti-mouse, 488 goat anti-rabbit, 694 goat anti-rat, and 488 goat anti-mouse) for 1 hour at room temperature.

#### Lamin B1 quantification

Quantification was done according to Chinta et al. (17). Z stacks images were taken using a confocal microscope. Once GFAP-positive cells had been identified, we searched for the z-section with the greatest Lamin B1 intensity. Lamin B1 intensity was measured by Image-J.

#### Colocalization analysis

Colocalization analysis was done with the imaris colocalization algorithm, which analyzes 2-channel confocal z-stacks. It measures the intensity of each channel voxel-by-voxel. The analysis of colocalization of astrocytes and p15ink4b proximal to the plaques was performed in an area starting from the plaque’s center up to 50 μm from its maximal width. The analysis of colocalization of astrocytes and p15ink4b distal to the plaques was performed in a 250×250 μm area free of plaques.

#### Isolation and treatment of adult astrocytes

Astrocytes were isolated and cultured as previously described by our group (11). In brief, the cerebellum and brain stem were removed from all brains. The cells were detached and passaged with 0.05% trypsin-ethylene diamine tetraacetic acid (Gibco) once on day 6–7 *in vitro* with a second and final passage on day 8–10. Before the second passage, the plates were placed on a shaking rotator (120 rotations per minute) for 2 hours at 37°C to remove all other cells. Cell purity was assessed, and subsequent experiments were conducted on day 11–15 *in vitro*. All experiments were done when the culture was confluent. The TGF-β1 inhibitor that was used is a selective inhibitor of ALK5, a receptor for TGF-β1, which phosphorylates Smad2 and Smad3 (18) (Selleckchem, # S1067). Aβ42 was purchased from GL Biochem, #052487. Pyrrolidinedithiocarbamic acid ammonium salt (PDTC) was used as an NF-κB inhibitor (#ALX-400-002). PDTC prevents the mobilization of NF-κB (19), and it had been shown to inhibit NF-κB in rat astrocytes (20).

#### Real-time polymerase chain reaction (RT-PCR)

RNA was extracted using the MasterPure™ RNA Purification Kit (Epicentre, #MC85200). Cells were analyzed for messenger RNA (mRNA) expression by reverse transcription (QUANTA Biosciences qscript, #95047) followed by RT-PCR with SYBR green (QUANTA Biosciences, #095073). Glyceraldehyde 3-phosphate dehydrogenase (GAPDH), a housekeeping gene, was used to normalize the expression of each gene in each sample.

#### Enzyme-linked immunosorbent assay (ELISA)

The active fragment of Aβ alone was shown not to result in significant IL-6 secretion in rat astrocytes (21). In order to induce cytokine release, the astrocyte cultures were stimulated using 5 μg/ml polyinosinic: polycytidylic βcid (Poly I:C, Sigma, #P4937) for 24 hours, after which the medium was collected (22). IL-6 and MIP-2 were measured using paired antibodies and recombinant cytokines according to the manufacturer’s recommendations (BD PharMingen Systems) (11).

#### Western Blot (WB)

WB was performed as described by Trudler et al. (23). Membranes were scanned and protein bands were normalized to GAPDH (Millipore, #MAB374). The relative band intensity was determined using Image-J. The antibodies used were as follows: pSMAD2 (1:000, Cell Signaling, #C-3108S), SMAD2 (1:1000, Cell Signaling, #C-3103S), p21 (1:1000, Santa Cruz, #sc-6246), vascular endothelial growth factor (VEGF) (Abcam, #ab46154), p65 (1:1000, Santa Cruz, #sc-372), brain-derived neurotrophic factor (BDNF) (1:2000, Abcam, #ab108319), and p15 (1:500, Assay Biotechnology, #C0287).

#### SA-β-gal for fluorescence-activated cell sorting (FACS)

FACS was performed according to Debacq-Chainiaux et al. (24). Briefly, in order to induce lysosomal alkalinization, subconfluent cells were pre-treated with 100 nM bafilomycin A1 for 1 hour, and C_12_FDG was then added to the medium to a final concentration of 30 μM for 2 hours. Next, the cells were harvested, and fluorescence was measured by FACS, after which the mean fluorescence intensity of the population was analyzed.

#### Neonatal neuronal culture

Neuronal isolation was performed as described in detail elsewhere (11). Briefly, cortices from 5 P1–2 BALB/c pups were dissected and placed on ice, chopped with scissors in a papain-based dissociation buffer [2.5 mM CaCl2, 0.83 mM EDTA, 137 U papain (Sigma), 100 μl DNAse (Sigma), 3–5 crystals of Cysin HBSS-HEPES (20 mM)], placed on a rotating shaker for 20 minutes at room temperature, and then centrifuged for 5 minutes at 1300 rpm. The pellet was resuspended with growth medium (5% fetal calf serum, 2% B27, 1% L-Gln, 0.5% PenStrep in Neurobasal A medium). The cells were placed in a 10-mm dish for 1 hour at 37°C, and the supernatant containing neurons was collected and used for co-culture experiments. The length of the longest neurite per neuron, as evidenced by β-tubulin staining, was measured in at least 100 neurons per coverslip.

#### Statistical Analysis

Results were analyzed using either two-tailed Student’s t-test when 2 groups were compared, and with a one-way analysis of variance followed by Bonferroni’s test when 3 or more groups were analyzed or planned comparisons were conducted. Differences were taken as being statistically significant at \**P* < 0.05, \*\**P* < 0.01, and \*\*\**P* < 0.001.

## RESULTS

### Changes in CS markers in 5xFAD astrocytes surrounding Aβ plaques

To investigate the regional link between the development of Aβ pathology and AS, we stained brain sections of 4- and 6-month-old WT and 5xFAD mice for the CS marker, p15ink4b (2), Aβ, and astrocytes activation (GFAP). We identified low levels of p15ink4b in both the 4-month-old WT and 5xFAD mice, who were at an earlier stage of Aβ plaque formation (Fig. 1A 1-8). At 6 months of age, we found significantly stronger staining of p15ink4b in astrocytes co-localized with Aβ plaques compared to distal astrocytes from Aβ plaques in the same brain region (hippocampus) and to WT mice (**Fig. 1A** **9-16**, **Fig. 1B**). These results suggest that senescent astrocytes accumulate at the sites of pathology. Reduction in lamin B1 levels was reported as being a marker of CS (25, 26). We discovered a significant reduction in lamin B1 in astrocytes in the 6-month-old 5xFAD mice compared to age-matched controls (**Fig. 2**, *P* = 0.02). Overall, these results indicate that 5xFAD astrocytes show markers of CS in 6-month-old mice in parallel with an increase in Aβ plaque load.

**Figure 1.**
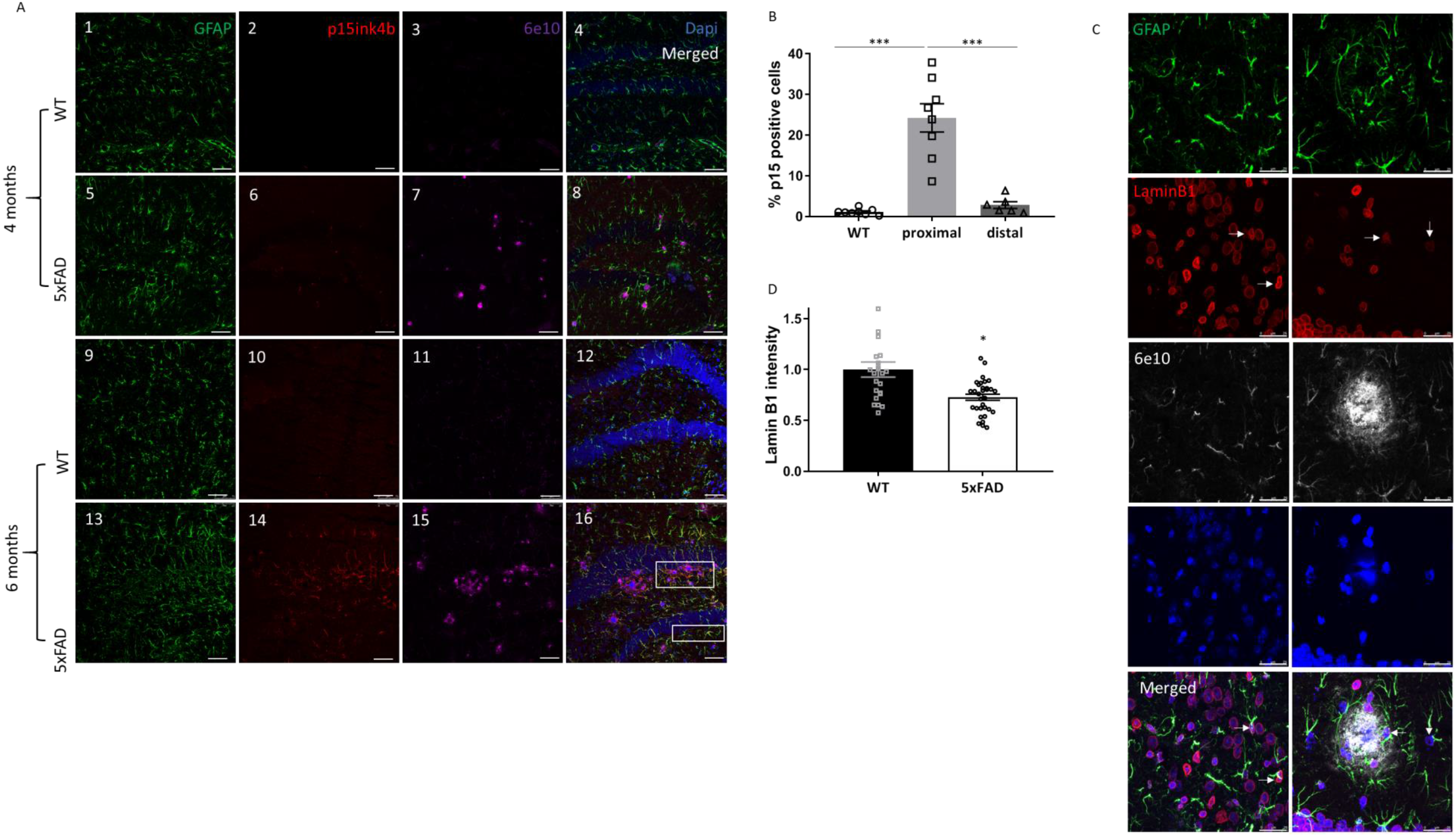
Changes in cellular senescence (CS) markers in 5xFAD astrocytes surrounding Aβ plaques. (A) Brain slices from 4- and 6-month-old wild-type (WT) and 5xFAD mice were stained for astrocytes (using GFAP antibody), CS (using p15 antibody), Aβ plaques (using 6e10 antibody), and cell nuclei (using 4’,6-diamidino-2-phenylindole; DAPI). Images 1–4: 4-month-old WT mice. Images 5-8: 4-month-old 5xFAD mice. Images 9–12: 6-month-old WT mice. Images 13-16: 6-month-old 5xFAD mice. Scale bar = 75 μm. n=8 (WT and 5xFAD proximal) and n=6 (5xFAD distal), with 2 mice in each group. (B) Quantification of co-localization between p15ink4b and GFAP. A planned comparisons test was conducted. (C) Brain slices from 6-month-old WT and 5xFAD mice were stained for astrocytes (using GFAP antibody), Lamin B1, Aβ plaques (using 6e10 antibody), and cell nuclei (using 4’,6-diamidino-2-phenylindole, DAPI). Arrows point to astrocyte nuclei. Scale bar=25 μM. (D) Quantification of Lamin B1 fluorescence intensity (n=23 astrocytes, 2 mice for the WT group and n=32 astrocytes, 2 mice for the 5xFAD group).

**Figure 2.**
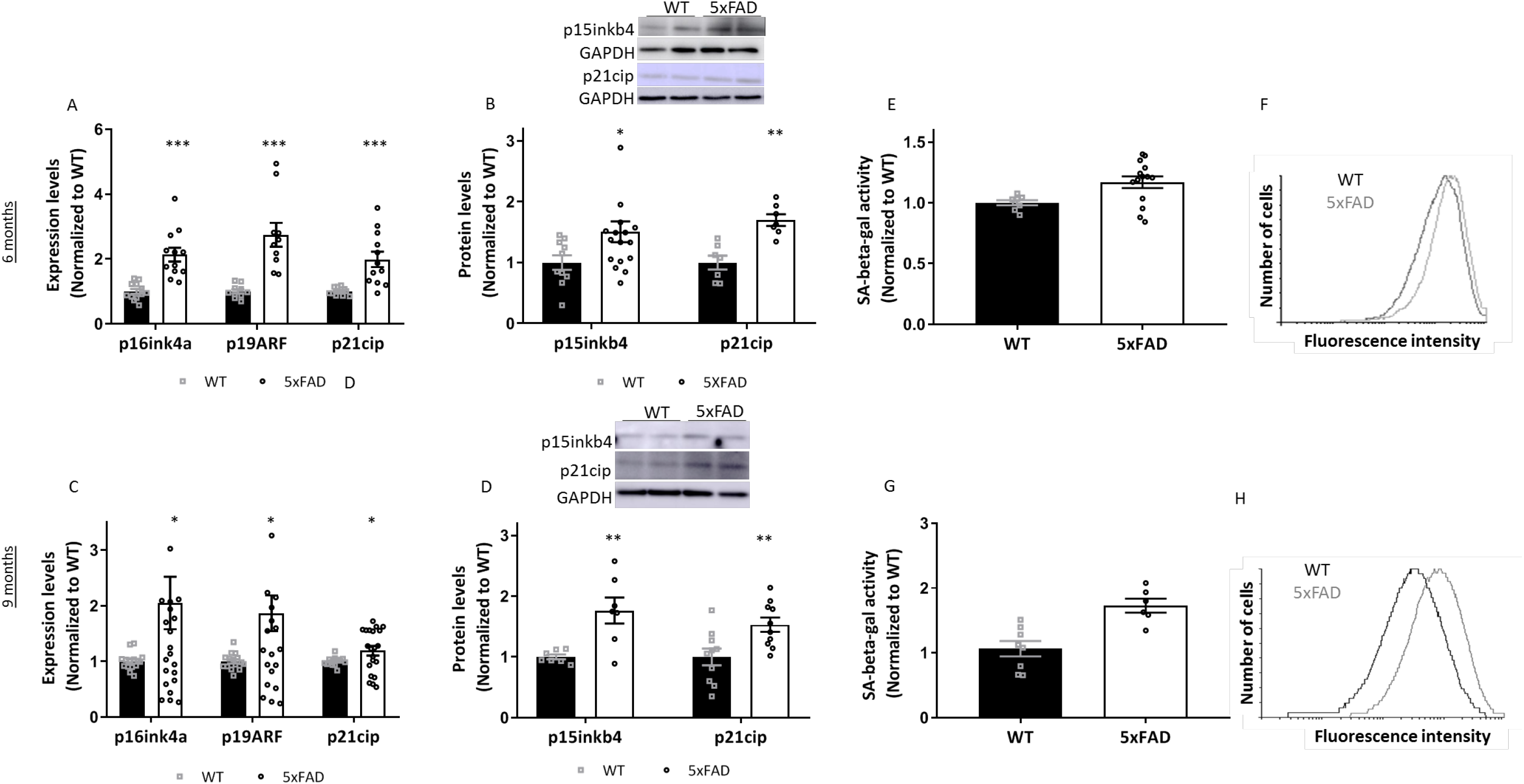
Increase in CS markers in isolated 5xFAD astrocytes. (A) p16^ink4a^ (n=12), p19^ARF^ (n=10), p21^Cip1^ (n=11) mRNA expression levels in astrocytes isolated from 6-month-old WT and 5xFAD mice. Independent, two-tailed, Student’s t-tests were conducted. (B) p15^inkb4^ [n=10 (WT), n=17 (5xFAD)] p21^cip1^ (n=10 for the WT group, n=13 for the 5xFAD group) protein levels were tested in astrocytes isolated from 6-month-old WT and 5xFAD mice. An independent, two-tailed Student’s t-test was conducted (n=7). (C) 6-month-old WT and 5xFAD astrocytes were tested for SA-β-gal activity using FACS. An independent, two-tailed Student’s t-test was conducted. [n=8 (WT), n=14 (5xFAD)]. (D) A representative graph of an SA-β-gal experiment. Astrocytes were isolated from 9-month-old WT and 5xFAD mice, and cells were tested for (E) mRNA expression of p16^ink4a^ [n=15 (WT), n=23 (5xFAD)], p19^ARF^ [n=15 (WT), n=22 (5xFAD)], p21^Cip1^ [n=15 (WT), n=21 (5xFAD)]; protein levels of p15 ^ink4b^ (n=7) and p21 ^Cip1^ (n=10) levels using WB. (F) SA-β-gal using FACS. An independent, two-tailed Student’s t-test was conducted [n=8 (WT) and n=10 (5xFAD)]. (G) A representative graph of an SA-β-gal experiment.

### Increase in CS markers in isolated 5xFAD astrocytes

We next assessed CS markers in astrocytes isolated from 5xFAD mice. We had recently reported a new protocol to isolate adult astrocytes from mice (11). Cultured 5xFAD astrocytes exhibited similar characteristics to *in* vivo astrocytes, such as deficits in the uptake of soluble Aβ and an increase in glial fibrillary acidic protein (GFAP) compared to WT astrocytes (11). We now isolated astrocytes from 6-month-old WT and 5xFAD mice and tested the expression of various CS markers. We found increased mRNA expression levels of CS markers, such as p16^ink4a^, p19^ARF^, and p21^Cip1^ (**Fig. 2A**, *P* < 0.001, *P* = 0.0001, and *P* = 0.0007, respectively). We also detected increased protein levels of 2 CS markers, p15^ink4b^ and p21^Cip1^ (**Fig. 2B**, *P* = 0.046 and *P* = 0.01, respectively) in 6-month-old 5xFAD astrocytes. CS markers were also observed in 5xFAD astrocytes isolated from 9-month-old mice. We found higher expression levels of p16^ink4a^, p19^ARF^, and p21^Cip1^ (**Fig. 2C**, *P* = 0.04, *P* = 0.017 and *P* = 0.03, respectively) as well as higher p15^inkb4^ and p21^Cip1^ protein levels (**Fig. 2D**, *P*= 0.002 and *P* = 0.004, respectively) in 5xFAD astrocytes compared to 9-month-old WT astrocytes. Senescence-associated β-gal was used as a marker of CS (27), and it was reportedly increased in senescent astrocytes (14, 28). To further characterize CS in astrocytes, we used FACS and assessed SA-β-gal activity. Our results revealed an increase in SA-β-gal activity in both 6- (**Fig. 2E, F**, *P* = 0.02) and 9-month-old (**Fig. 2G H**, P < 0.001) 5xFAD astrocytes compared to age-matched controls. These findings suggest that an increase in CS marker expression in 5xFAD astrocytes parallels chronic exposure to Aβ pathology.

### Impairment of 5xFAD astrocytes in supporting neuronal growth in an NF-κB-dependent manner

The NF-κB pathway was suggested to play an important role in mediating SASP components (12). Using a co-culture of neurons and 5XFAD astrocytes, we now observed that neurons grown on 9-month-old 5xFAD astrocytes have shorter neurites compared to neurons grown on age-matched WT astrocytes (**Fig. 3A-B**, *P* = 0.03). This finding was NF-κB-dependent, since an inhibition of NF-κB resulted in an increase in neurite length in neurons grown on 5xFAD astrocytes (**Fig. 3A, B**, *P* = 0.02). We discovered that inhibition of NF-κB reduced IL-6 levels (**Fig. 3C**, Group effect: *P* < 0.001), which we had previously suggested to play a role in 5xFAD astrocyte neurotoxicity (Iram et al. 2016). There was also a decrease in the expression of VEGF and BDNF proteins in 9-month-old 5xFAD astrocytes (**Fig. 3D**, VEGF: *P* = 0.0005; BDNF: *P* = 0.0001). These results suggest that senescent astrocytes fail to support neurons and that this failure can be reversed by inhibiting the NF-kB pathway.

**Figure 3.**
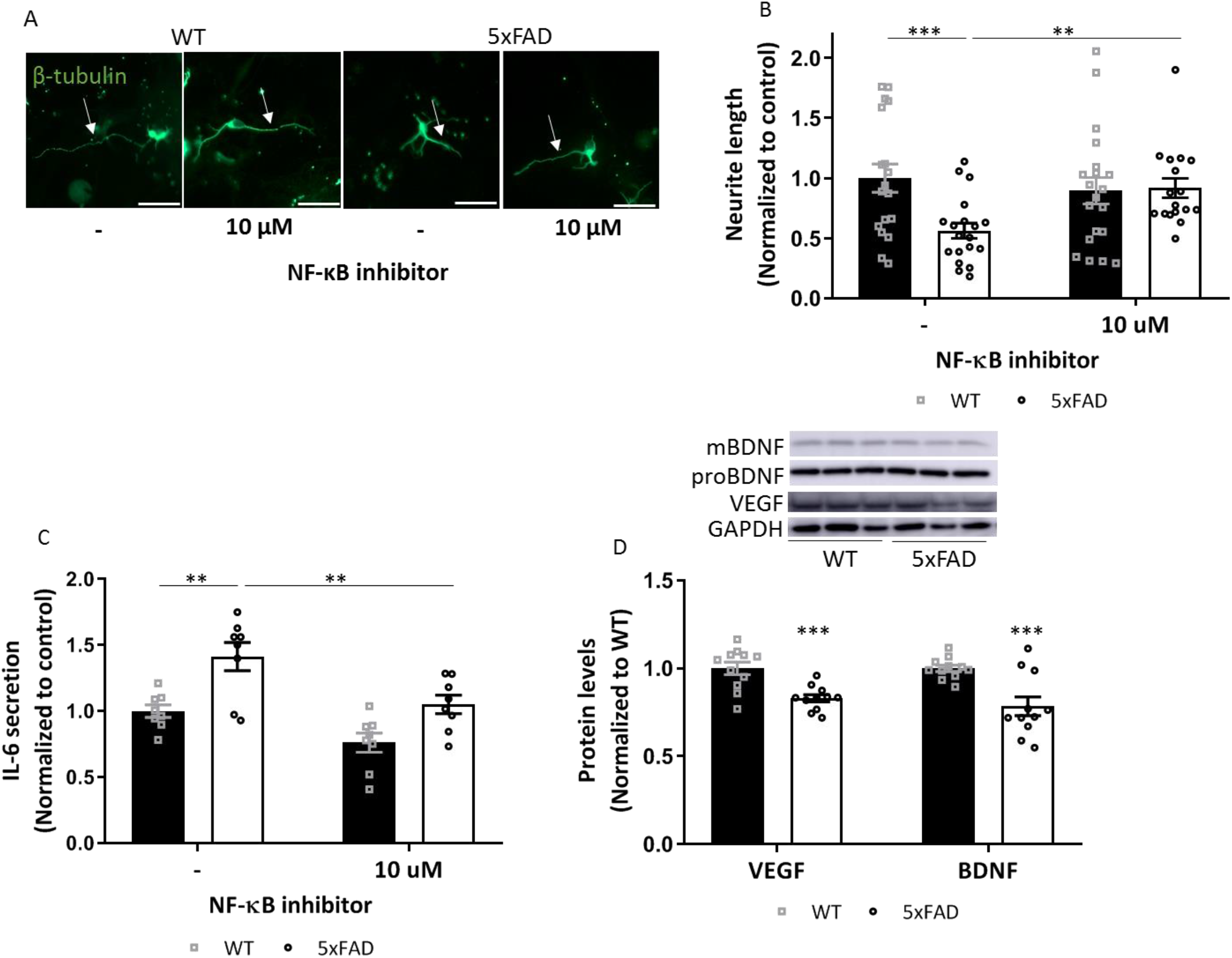
Impairment of 5xFAD astrocytes in supporting neuronal growth in an NF-κB-dependent manner. (A) Representative images of co-cultures of neurons and astrocytes stained for β-tubulin. (B) Neurite length of neurons grown on a monolayer of astrocytes isolated from 9-month-old wild-type (WT) and 5xFAD mice. Astrocytes were pre-incubated overnight with NF-κB inhibitor. (n=25 neurons, 4 mice in each group). Scale bar = 25 μM. (C) Astrocytes from 9-month-old WT and 5xFAD mice were incubated overnight with 10 μM NF-κB inhibitor and 5 μg/ml poly I:C, and IL-6 secretion levels were tested using ELISA (n=8). (D) BDNF and VEGF protein levels in astrocytes from 9-month-old WT and 5xFAD mice. An independent, two-tailed Student’s t-test was conducted (n=11).

### AS is mediated by the TGF-β1-SMAD2/3 pathway

TGF-β1 was suggested as being secreted as part of the SASP and to induce CS (29). We had earlier reported an increase in TGF-β1 expression in the brains of 5xFAD mice (30). We now demonstrated that TGF-β1 expression is increased in 6-month-old 5xFAD astrocytes (**Fig. 4A**, *P* < 0.001). We also observed an increase in TGF-β1-SMAD2/3 pathway induction in 5xFAD astrocytes compared to WT astrocytes (**Fig. 4B**, *P* = 0.0008). To further investigate the role of TGF-β1 in mediating astrocyte senescence, we incubated astrocytes isolated from 3-month-old WT mice with TGF-β1 and found a significant increase in p16^ink4a^ and p21^Cip1^ expression levels (**Fig. 4C**, *P* = 0.01 and *P* = 0.0007, respectively). To further assess the role of TGF-β1 expression in astrocytes, we isolated astrocytes from WT and GFAP-TGF-β1 mice that overexpress TGF-β1 under the astrocyte promoter, GFAP (15). The results were a significant increase in CS expression markers, such as p16^ink4a^, p19^ARF^ and p21^Cip1^, already present in astrocytes isolated from 3-month-old TGF-β1 mice (**Fig. 4D**, *P* = 0.001, *P* = 0.0006, and *P* =0.003, respectively), as well as an increase in SA-β-gal activity in TGF-β1 astrocytes **(Fig. 4E F**, *P* = 0.0001). We further assessed the SASP profile of astrocytes and observed an increase in both macrophage inflammatory protein 2 (MIP-2) and IL-6 secretion by TGF-β1 astrocytes (**Fig. 4G**, *P* = 0.01 and *P* = 0.02, respectively). To assess whether AS is TGF-β1-dependent, we incubated TGF-β1 astrocytes with a TGF-β1-activin receptor-like kinase (ALK)5 inhibitor (hereafter, TGF-β1 inhibitor) and observed a reduction in p16^ink4a^ levels compared to the control group (**Fig. 4H**, *P* < 0.001). It also emerged that inhibition of the TGF-β1-SMAD2/3 pathway in 5xFAD astrocytes resulted in a reduction of both the expression levels of p16^ink4a^ (**Fig. 4I**, *P* = 0.03) and the secretion of the pro-inflammatory cytokine IL-6 (**Fig. 4J**, *P* = 0.003). These results suggest that TGF-β1 can mediate 5xFAD in AS.

**Figure 4.**
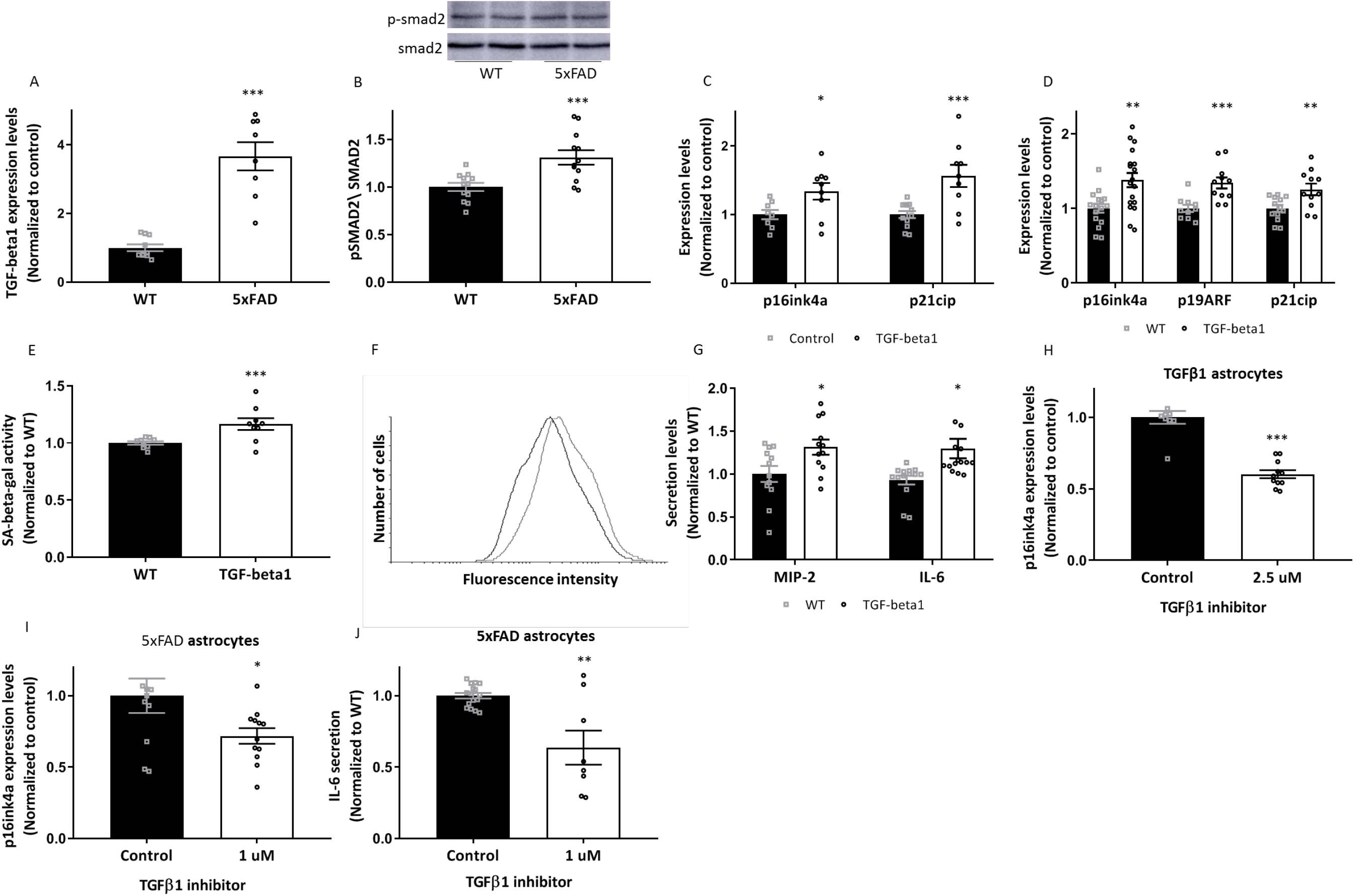
AS is mediated by the TGF-β1-SMAD2/3 pathway. (A) TGF-β1 expression levels in astrocytes isolated from 6-month-old wild-type (WT) and 5xFAD mice. An independent, two-tailed Student’s t-test was conducted (n=8). (B) SMAD2 phosphorylation levels were tested and normalized to total SMAD2 levels in astrocytes isolated from 6-month-old WT and 5xFAD mice. An independent, two-tailed Student’s t-test was conducted (n=12). (C) WT astrocytes were incubated with 5 ng/ml of TGF-β1 for 6 hours, and p16^ink4a^ and p21^Cip1^ expression levels were measured. Independent, two-tailed Student’s t-tests were conducted (n=9). (D) p16^ink4a^ (n=16), p19^ARF^ (n=10) and p21^Cip1^ (n=13) mRNA levels in astrocytes isolated from WT and TGF-β1 mice. Independent, two-tailed Student’s t-tests were conducted. (E) SA-β-gal activity was measured in astrocytes isolated from WT and TGF-β1 mice using FACS. An independent, two-tailed Student’s t-test was conducted (n=9). (F) A representative graph of an SA-β-gal experiment. (G) Astrocytes isolated from WT and TGF-β1 mice were incubated with 5 μg/ml of poly I:C for overnight, and IL-6 (n=15) and MIP-2 (n=12) secretion levels were measured. Independent, two-tailed Student’s t-tests were conducted. (H) Astrocytes from TGF-β1 mice were incubated with TGF-β1-SMAD2/3 inhibitor for 6 hours and then tested for p16^ink4a^ expression levels. An independent, two-tailed Student’s t-test was conducted (n=9). (I) Astrocytes isolated from 5xFAD mice were incubated with 1 μM TGF-β1-SMAD2/3 inhibitor for 6 hours and tested for expression levels of p16^ink4a^. An independent, two-tailed Student’s t-test was conducted (n=11). (J) 5xFAD astrocytes were incubated with 1 μM TGF-β1-SMAD2/3 inhibitor and 5 μg/ml of poly I:C for ON and the supernatant was tested for IL-6 secretion. An independent, two-tailed Student’s t-test was conducted (n=15).

### Aβ42 induces AS through the TGF-β1-SMAD2/3-pathway

It has been suggested that Aβ1-42 induces AS (14). We examined whether this effect is mediated by the TGF-β1-SMAD2/3 pathway. For this purpose, we incubated young WT astrocytes with Aβ1-42, which led to a significant increase in TGF-β1 SMAD2/3 phosphorylation (**Fig. 5A**, *P* = 0.005) as well as in p21^Cip1^ levels (**Fig. 5B**, *P* = 0.003). Inhibition of the TGF-β1-SMAD2/3 pathway also prevented AS following incubation with Aβ1-42 (**Fig. 5B**, P = 0.03).

**Figure 5.**
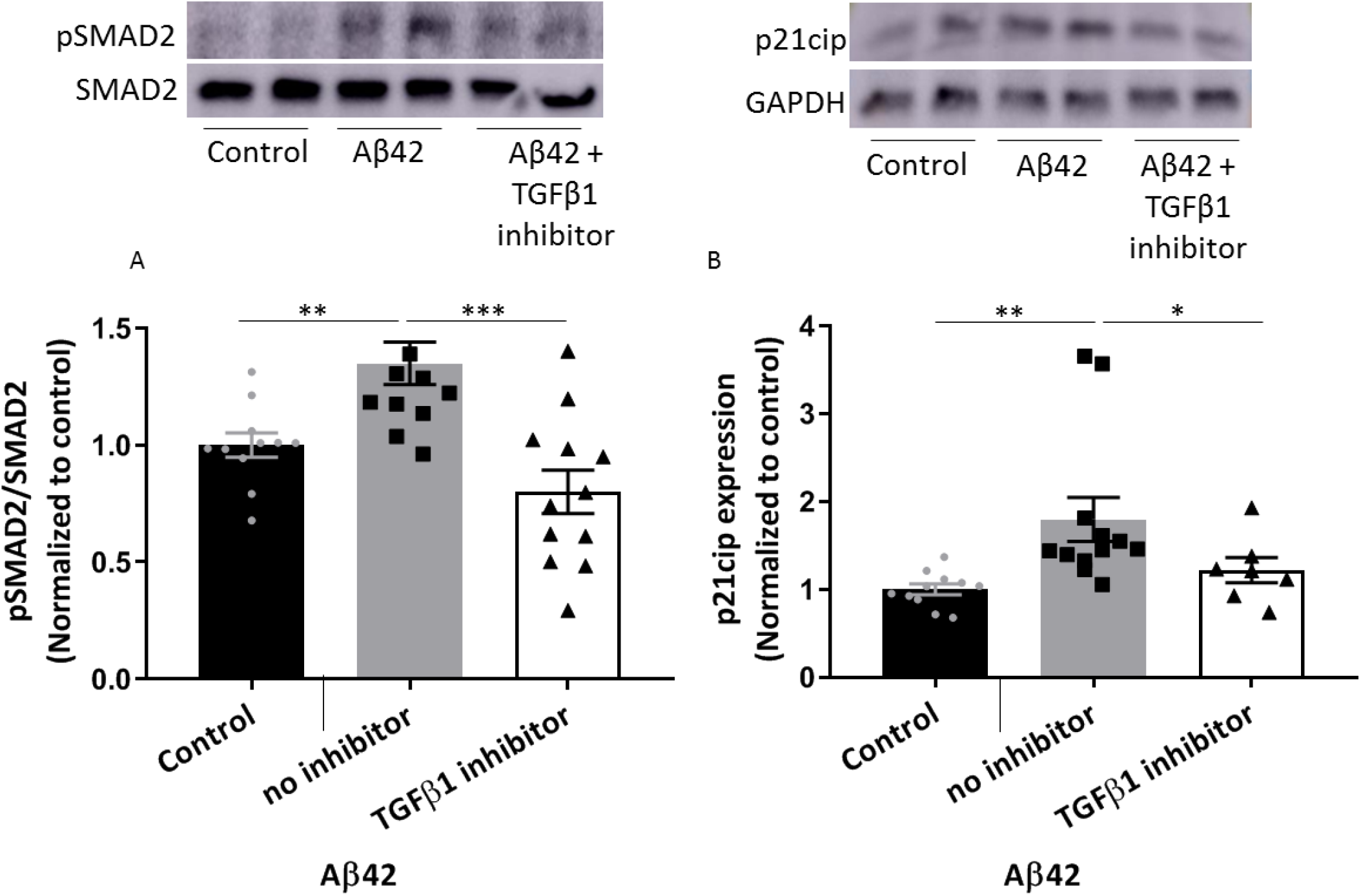
Aβ42 induces astrocyte senescence through the TGF-β1-SMAD2/3 pathway. Wild-type (WT) astrocytes were incubated with 10 μM of soluble Aβ with or without 5 μM of a TGF-β1-SMAD2/3 inhibitor for 24 hours and then (A) measured for the induction of the SMAD2/3 pathway (n=12), and (B) p21^Cip1^ levels (n=9). Planned comparisons were conducted.

## DISCUSSION

The present study aimed to test our hypothesis that with age, astrocytes in 5xFAD mice would exhibit a CS phenotype that could promote neurodegeneration. We discovered that an increase in Aβ plaque load may lead to CS in astrocytes, leading us to suggest that this outcome is mediated through the activation of the TGF-β1 pathway. It emerged that senescent astrocytes fail to support neurons, so that an inhibition of the NF-κB pathway may improve the astrocytes’ neuroprotective profile.

Given the prominent role of astrocytes in a wide variety of brain functions, AS is likely to have implications on the disease course of AD as well. Astrocytes exhibited characteristics of CS and SASP in AD (14), in a model of amyotrophic lateral sclerosis (31), in Parkinson’s disease (17), and in tauopathy (9), suggesting that astrocytes have a predisposition to senescence. Here, we show that astrocytes surrounding Aβ plaques exhibited markers of CS in an age-dependent manner.

Astrocytes were reported to be prone to stress-induced senescence, such as oxidative stress (32) and Aβ1-42 (14). Using astrocyte-neuron co-cultures, we now showed that 5xFAD astrocytes fail to support neurons compared to WT astrocytes. Overall, these results indicate that AS may have a physiological relevance to the disease.

Although senescence is considered to be a major part of the aging process and in age-related diseases (3), little is known about the specific mechanisms leading cells to enter senescence. TGF-β1 levels were increased with aging (33), and in the ante-and post-mortem sera from patients with AD, as well as in the postmortem cerebrospinal fluid of patients with AD (34). Furthermore, TGF-β is secreted as part of the SASP (3), and it can trigger senescence in neighboring cells (3, 35, 36). Our current study findings demonstrated that the TGF-β1-SMAD2/3 pathway is induced in 5xFAD astrocytes, and that inhibition of this pathway can reduce the expression of the senescence phenotype. Bhat et al.’s in vitro study provided evidence that Aβ42 leads to AS (14). Here, we show that Aβ42 can induce senescence in young astrocytes through an SMAD2/3-dependent pathway. Our results are in line with those of van der Wal et al. who reported that TGF-β1 accumulates around Aβ plaques (37) and, specifically, in astrocytes around Aβ plaques (38). These findings suggest that Aβ1-42 accumulates in the brain of AD patients, and that this accumulation upregulates the TGF-β1 SMAD2/3 pathway in astrocytes, thus leading to the induction of AS.

IL-6 is one of the prominent cytokines of the SASP (39), and overexpression of IL-6 in astrocytes can lead to the development of a neurologic disease characterized by seizures, ataxia, and tremors (40). Moreover, in 5xFAD mice, IL-6 was correlated with cognitive changes (41). We found that the failure of astrocytes from 9-month-old 5xFAD mice to support neurons was recovered by NF-κB inhibition. Furthermore, we observed an increase in IL-6 secretion in astrocytes from 9-month-old 5xFAD mice that was also mediated by NF-κB. We suggest that IL-6 in combination with the reduction in VEGF and BDNF plays a part in the impairment of 5xFAD astrocytes to support neurons. In line with this concept, Turnquist et al.’s study showed that while young astrocytes (early passage) displayed an increase in neuroprotective growth factors, which resulted in normal neurons, there was an increase in IL-6 secretion in senescent astrocytes (late passage) followed by apoptosis of neurons (42). Those authors also reported that brain sections from deceased patients with AD had increased numbers of senescent astrocytes as well as an increase in IL-6 expression (42).

Not only astrocytes change during aging and in neurodegenerative diseases. It has been suggested that microglia from old mice are less ramified compared to those from young mice, and that this process is accelerated in AD mice (43). Moreover, studies have shown that microglia cultured from aging brains (44) and AD brains (45) have increased production of pro-inflammatory cytokines, such as IL-6. While we suggest a role for senescent astrocytes in 5XFAD mice brain pathology, another report suggests the involvement of senescence in other types of glia cells in AD pathology. Bussian et al. (9) showed that microglia and astrocytes exhibit markers of CS in tauopathy, and that clearance of those senescent cells can ameliorate the disease course (9). It has also been reported in another AD mouse model that the increase of CS in NG2-expressing oligodendrocyte progenitor cells is linked to aggregating Aβ (46). Overall, these papers suggest an important role of senescent glia in neurodegenerative diseases.

Our findings indicate that there is an increase in AS in 5xFAD mice, and that Aβ induces this process through TGF-β1 activation. Our data demonstrate that senescence can lead to toxicity in neighboring cells. Understanding the role of AS in AD and the mechanisms that lead to AS may provide further insights into the role of astrocytes in the etiology of AD and other neurodegenerative diseases, and may aid in the identification of new targets for therapeutic intervention.

## ABBREVIATIONS

AD: Alzheimer’s disease
Aβ: amyloid-β
SASP: senescence-associated secretory phenotype
CS: cellular senescence
AS: astrocyte senescence
NF-κB: kappa-light-chain-enhancer of activated B cells
IL-6: interleuken-6
TGF β1: transforming growth factor β
PBS: phosphate buffered saline
PDTC: pyrrolidinedithiocarbamic acid ammonium salt
GAPDH: glyceraldehyde 3-phosphate dehydrogenase
WT: wild-type
(RT-PCR): Real-time polymerase chain reaction
mRNA: messenger RNA
ELISA: enzyme-linked immunosorbent assay
WB: western blot
NF-κB: nuclear factor kappa-light-chain-enhancer of activated B cells
VEGF: vascular endothelial growth factor
BDNF: brain-derived neurotrophic factor
GFAP: glial fibrillary acidic protein

## Acknowledgements

We thank Profs. Ari Barzilai and Yoav Henis from Tel Aviv University for helpful advice. We thank Ronit Galron for the assistance in the preparation of the co-culture and the WB. We thank Prof. Tony Wyss-Coray for the TGF-β1 mice.

## Author contributions

Amram S. and Frenkel D. conceived and planned the experiments, Amram S. executed all the experiments, and Amram S. and Frenkel D. analyzed the data and wrote the manuscript. Iram T. assisted with co-culture experiments and with the isolation protocol of astrocytes. Lazdon E. assisted with the WB experiments. Vasser R. and Ben Porath I. for crucial help for paper writing.

## Funding

This study was supported by grants from the JPco-fuND-European research for neurodegenerative disease D.F. PROP-AD is an EU Joint Programme - Neurodegenerative Disease Research (JPND) project. The project is supported through the following funding organisations under the aegis of JPND - www.jpnd.eu (AKA #301228 - Finland, BMBF #01ED1605 - Germany, CSO-MOH #30000-12631 - Israel, NFR #260786 - Norway, SRC #2015-06795 - Sweden). This project has received funding from the European Union’s Horizon 2020 research and innovation programme under grant agreement #643417 (JPco-fuND).

## Competing interests

None to report.

